# EPI Distortion Correction is Easy and Useful, and You Should Use It: A case study with toddler data

**DOI:** 10.1101/2020.09.28.306787

**Authors:** Vinai Roopchansingh, Jerry J. French, Dylan M. Nielson, Richard C. Reynolds, Daniel R. Glen, Precilla D’ Souza, Paul A. Taylor, Robert W. Cox, Audrey E. Thurm

## Abstract

Task, resting state, and diffusion MRI data are usually acquired from subjects using echo-planar based imaging techniques. These techniques are highly susceptible to B_0_ homogeneity effects that result in geometric distortions in the reconstructed images. As researchers work to link the information from these scans back to various developmental stages, or to conditions and diseases in specific regions or structures of the brain, it becomes critical to have accurate correspondence between more geometrically distorted echo-planar images and less geometrically distorted anatomical images. A variety of techniques and tools have been developed to improve this correspondence. The basic premise behind most techniques used to mitigate geometric distortion is to acquire enough information to inform software tools how echo-planar images are warped, then have them undo that warping. Here, we investigate the application of two common methods: B_0_ correction, and reverse-polarity phase-encoding (or reverse blip) correction. We implement each of these in two separate, widely used software packages in the field: AFNI and FSL. We find that using either technique in either software package results in reduced geometric distortions in the EPI images. We discuss the practical implementations of these methods (e.g., increased scan and processing time). In general, however, both methods possess readily available data acquisition schemes, and are highly efficient to include in processing streams. Due to the overall data improvement, we strongly recommend that researchers include one of these methods in their standard protocols.

## Introduction

The use of task and resting state EPI data to characterize the full range of both normal and pathological conditions [1] in various populations continues to increase. As use of such data increases, so does the need to make sure results obtained from it align more accurately and consistently with corresponding anatomical data, both within and across subjects, to maximize the validity of findings gleaned from studies utilizing EPI-based scanning techniques.

The toddlers whose data is used in the present methodological investigation were enrolled in a separate study of language delay. The group is comprised of both typically developing toddlers, and those with a familial risk of language delay. All participants underwent a scanning battery in which several types of image data were repeatedly acquired over multiple visits. These data include T_1_-weighted anatomical data, resting state EPI data, and auxiliary data, including, but not limited to, B_0_ field maps [2] and reversed-blip EPI data [3, 4].

In this work, we demonstrate how B_0_ and reversed-blip EPI data correct geometrically distorted EPI data and improve registration to T_1_-weighted structural data. This improvement is seen when using either of two separate, standard software suites (AFNI [5] and FSL [6]). The before-and-after comparisons are evaluated in two ways. First, we used ANTs [7] to quantify the global correspondence and alignment between the T_1_-weighted structural data and the EPIs, based on relative cost function values. Second, we visualized distortion and correction effects by comparing tissue segmentation boundaries estimated by FreeSurfer [8] from the T_1_-weighted structural data to corresponding features in uncorrected versus corrected EPI data. Finally, we discuss the practical implementations of each approach, both in terms of acquisition and analysis, and find that the uniform benefits of distortion correction greatly outweigh the minor increases in scanning and computational time.

## Methods

### Scanning

23 toddlers (age range: 14-45 months), enrolled in a study based on their status as typically developing (M=6, F=7) or due to identified language delays (M=7, F=3), were scanned on a General Electric (Waukesha, WI) MR-750 3T scanner (software revision DV22.0_V02), using the vendor’s 32-channel head coil. Participants were scanned unsedated, but asleep, during late-evening, running into early-morning sessions. Of the 70 scanning sessions that were performed on this group, 23 were acquired with full or partial data and included in these analyses. These scans were conducted with the approval of the NIH’s IRB, under ClinicalTrials.gov study ID NCT01339767.

For inclusion in this analysis, sessions must have a T_1_-weighted anatomical scan, an EPI scan with at least 15 volumes, and either a reversed-blip EPI scan; or a B_0_ scan. The included reversed-blip EPI or B_0_ scan must have been acquired with the subject’s head in the same position as the resting state data, so that at least one correction algorithm from each software package could be evaluated, and that the measured field and distortions would be matched to the EPI time-series data being collected. Of the subjects included in this analysis, there were 7 that had both a reversed-blip EPI scan and a B_0_ scan (in addition to the resting state and T_1_-weighted anatomical), which allowed for full cross-methodological comparison.

T_1_-weighted anatomical scans were acquired with the following parameters: image matrix = 256 × 256 × 142, voxel size of 0.75 × 0.75 × 1.0 mm^3^, TR = 2100 ms, TI = 885 ms, TE = 2.8 ms, acquisition BW = 50 kHz. Resting state data were acquired on a 96 × 96 image matrix, with 36 slices, TR = 2000 ms, TE = 30 ms, a nominal voxel size of 2.0 × 2.0 × 3.0 mm^3^, BW = 500 kHz, flip angle = 65°, EPI echo spacing = 620 µs, and an acceleration (SENSE) factor of 2. Reversed blip EPI data were collected matching parameters of the resting state scan, with the image phase-encoding blips having reversed polarity. B_0_ data were also acquired on the same image matrix and the same geometric prescription as the EPI data, using a dual-TE 2D gradient-echo sequence with the following parameters: TR = 1000 ms, TE1/TE2 = 6.4/8.86 ms, flip angle = 35°.

### Analysis

Tools in two common software packages (AFNI, version AFNI_19.2.23 ‘Claudius’; and FSL, version 6.0.1) were used to perform distortion correction on the EPI data using different algorithms and their accompanying software modules. From FSL, ‘topup’ and ‘applytopup’ were used to perform corrections using the reversed-blip EPI data, and ‘fugue’ was utilized with the B_0_ scan for distortion correction. From AFNI, the nonlinear warping tool ‘3dQwarp’ [9] leveraged information from the reversed-blip EPI scan data. This module is utilized in both ‘unWarpEPI.py’ (as a stand-alone program) and ‘afni_proc.py’ (as a portion of the overall data pre-processing and processing stream). A series of commands eventually employing ‘3dNwarpApply’ used B_0_ data for distortion correction. The procedure utilizing ‘3dNwarpApply’ for B_0_ corrections in AFNI is currently available in the ‘epi_b0_correct.py’ module. T_1_-weighted anatomical data were skull-stripped with ‘3dSkullStrip’ in AFNI.

The first 15 volumes of EPI data from each session were distortion corrected using either the B_0_ or reversed-blip EPI data, via the appropriate modules from each of the software suites. After distortion correction, the EPI data were resampled using cubic interpolation to the same grid as the T_1_-weighted anatomical data using AFNI’s ‘3dresample’. The ‘ImageMath’ module in ANTs was then used to compute the Pearson Correlation (PC) between the resampled distortion-corrected EPI time-series data, and the skull-stripped T_1_-weighted data. In some sessions, data were collected with the subjects’ heads in different positions between the anatomical and resting state data. This was most likely due to the toddler having to be removed from the MR scanner mid-session because they had woken up, were soothed, and put back into the MR scanner, asleep in a different position. For these cases, the ‘antsRegistration’ module was used to perform a rigid-body registration of the EPI data to the anatomical data as the final step in the pipeline, after distortion corrections and resampling to the anatomical data space. All results presented below include cases only where data were fully processed, and all stages (from distortion correction to rigid-body registration) were successfully completed.

A fractional change in Fisher-transformed Pearson Correlation (**Z-PC**) was then computed for each pipeline, with the score of the alignment of the uncorrected EPI data to the T_1_-weighted anatomical data serving as the base value from which the fractional change was computed. Z-PC changes for each pipeline were evaluated with a within-subject t-test. Comparisons between pipelines were made with within-subject tests when possible, and between-subject t-tests otherwise.

Given the geometrical nature of the EPI distortion, the performance of all software algorithm combinations were also evaluated visually, by comparing the location of structures in the anatomical volume with those in the uncorrected and corrected EPI data. This direct comparison provides a detailed examination of the volumes and local assessment of distortions. FreeSurfer’s (version 6.0.0) ‘recon-all’ tool was run on each subject’s T_1_-weighted anatomical volume, and edges between estimated tissue classes (gray matter, white matter and CSF combined with ventricles) were calculated in AFNI. These edges were overlaid on both the original and corrected volumes, for comparison of the relatively undistorted T_1_-weighted structures with those of the uncorrected and corrected EPI data. Locations of large distortion and/or relative difference in correction algorithms were noted and highlighted. To reduce the effects of brightness inhomogeneities present in EPI volumes, the EPIs presented in the figures were brightness-adjusted (“unifized”) to clarify tissue types and boundaries using AFNI’s ‘3dUnifize’ program with the “-EPI” option.

To facilitate replication of the various distortion correction streams on the EPI data, and comparison of the distortion-corrected EPI data to the anatomical data, data were organized into a BIDS tree structure [10]. The commands and parameters used in this study are available in a publicly accessible repository on GitHub.com (https://github.com/nih-fmrif/bids-b0-tools).

## Results

In all figures and tables below, the following keys for software packages and distortion correction techniques are used: A - Correction done with AFNI’s tools; F - Correction done with FSL’s tools; B - B_0_-based algorithms utilized for correction; E - EPI reversed-blip algorithms used for correction. Thus, “AB” refers to B_0_-based correction using AFNI, etc. Additionally, “DOF” refers to the number of degrees of freedom in each test.

Figure 1 shows the change in Z-PC between EPI and anatomical datasets for each subject, before (“uncorrected”) and after (“corrected”) geometric correction of EPI data. In every case but one, there was an increase in Z-PC for the corrected EPI. This was from 10 subjects who had all of the data necessary to perform and evaluate the B_0_ correction pipelines, 22 subjects who had all of the data necessary to perform the reversed-blip EPI correction pipelines, and had successful registration of both uncorrected and corrected EPI data to the anatomical data. Thus, across both techniques and software packages, EPI distortion correction increased similarity to the anatomical volume. Table 1 shows the accompanying table of one-sample t-test of changes in the Z-PC scores, which were significant for all package and correction algorithm combinations.

**Table 1:** Comparisons of Z-PC scores of corrected EPI data to T_1_-weighted anatomical data, versus uncorrected EPI data to T_1_-weighted anatomical data, across techniques and software packages (see Fig. 1). In all cases the mean Z-PC scores of the corrected volumes were higher than for the uncorrected volumes.

| Package/Correction Algorithm | T-statistic (DOF) | Corrected *p*-value |
| --- | --- | --- |
| AB | 11.695 (9) | 3.83e-06 |
| AE | 5.885 (21) | 3.07e-05 |
| FB | 10.728 (9) | 7.95e-06 |
| FE | 5.150 (21) | 1.68e-04 |

**Table 2:** Z-PC score comparison between each software packages and correction technique combination. No significant mean Z-PC difference was estimated between any pair.

| Package/Algorithm Comparison | T-statistic (DOF) | Corrected *p*-values |
| --- | --- | --- |
| AB vs. AE | 0.343 (9) | 1.0 |
| AB vs. FB | -0.278 (9) | 1.0 |
| AB vs. FE | 0.663 (9) | 1.0 |
| AE vs. FB | -0.746 (9) | 1.0 |
| AE vs. FE | 0.484 (21) | 1.0 |
| FB vs. FE | 1.047 (9) | 1.0 |

**Figure 1:**
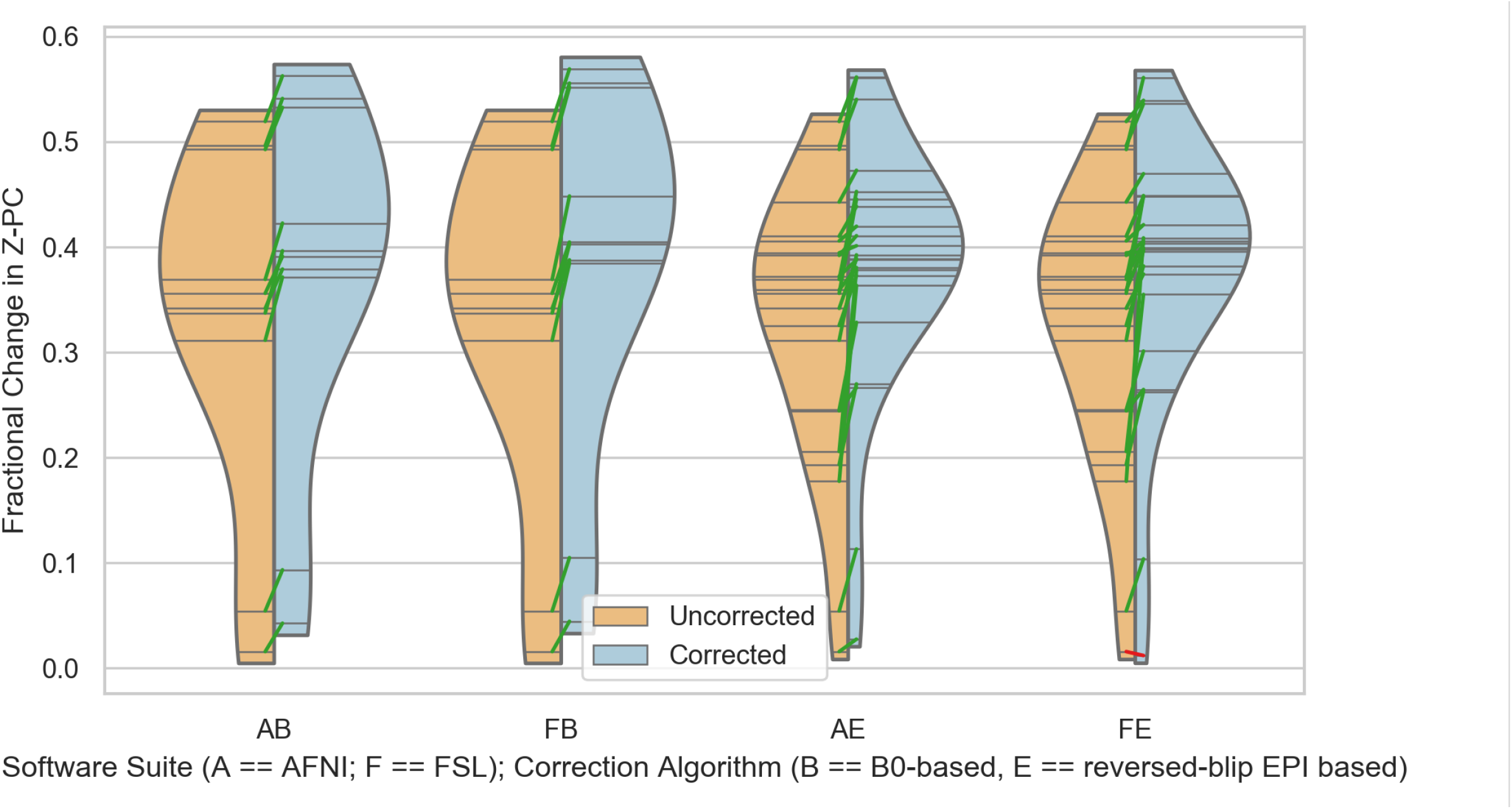
Changes in Z-PC for all software packages and algorithms, between uncorrected (orange) and corrected (blue) EPI data. N=10 for B_0_-based correction, and N=22 for reversed-blip correction.

When the performances of the packages were compared against each other, there was no significant difference in performance measured with Z-PC. This null result was likely due to the combination of the size of the improvement, in conjunction with the small number of subjects in each distortion correction test group. Furthermore, Z-PC is an omnibus measure across the full volume, and detailed comparisons would require local structural comparisons.

Figure 2 allows a direct visual comparison of distorted and corrected EPIs for each of the 7 subjects in the present study who completed the full set of scans (anatomical, EPI, B_0_, and reverse-blip). Each row shows the data from within the same location, within the same subject. The red lines denote tissues boundaries from FreeSurfer parcellation. Green and blue arrows highlight major distortions in the original EPI for a given subject. The improvement in correspondence between the structure of the final EPI and that of the anatomical volume can be observed for each software-technique combination. This overall improvement in structural overlap is also present in the results from the remaining subjects, who only completed scanning for one of the distortion correction methods.

**Figure 2:**
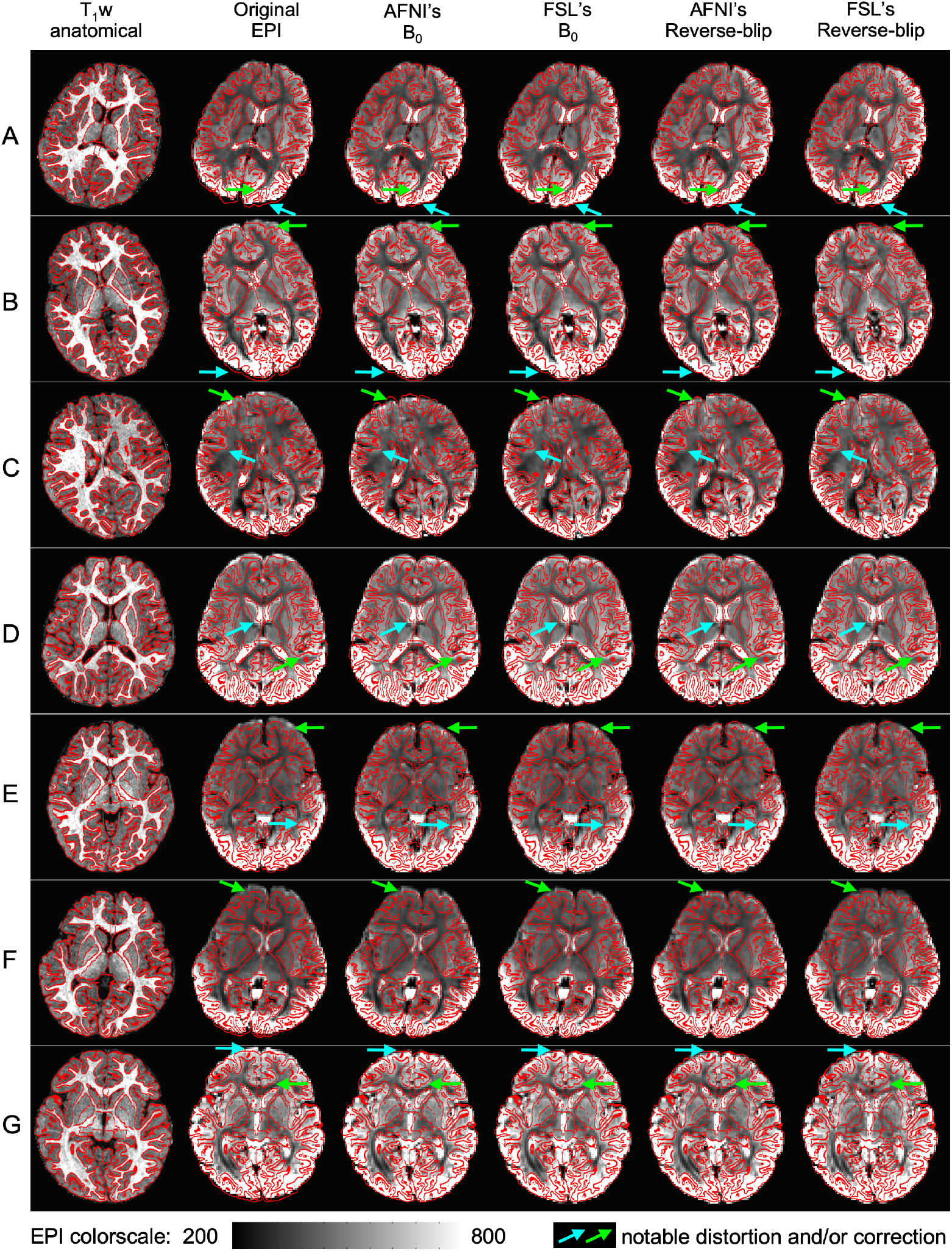
Axial slices (left=left) from each of the 7 subjects (per row, A-G) in the present study who completed all scans. Column labels describe volume type and processing. For each subject, tissue boundaries from FreeSurfer parcellation are overlaid (red lines). Additionally, green and blue arrows highlight locations of large distortion in the original EPI, and the relative correction of each approach can be assessed.

## Discussion and Conclusions

The purpose of this study was to evaluate the scale and effectiveness of EPI distortion correction in standard FMRI applications. Multiple techniques and software packages were implemented on the same datasets. Importantly, the results of both the quantitative and qualitative comparisons made here clearly show the benefit of performing EPI distortion correction, regardless of the algorithm or software suite used.

The two algorithms whose performance was explored here (B_0_-based and reverse blip-based) each have advantages and disadvantages, which can make them more or less suitable for any given study. We briefly describe the theoretical background of each algorithm, as well as practical aspects of using each.

For B_0_-based algorithms and correction techniques, the B_0_ field map is an actual measure of the main static magnetic field; this dataset is effectively a displacement map that is used to move image energy around, grounded in the physics of the B_0_ inhomogeneity. The reversed-blip approach is more empirical and based on image-processing: one acquires pairs of EPI volumes that should have “opposite” warps along the phase-encoding direction, and then applies volumetric alignment to “unwarp” the data from itself, and from this - estimates a warp field.

On a practical note, 2D gradient-echo based B_0_ data typically have a higher signal-to-noise ratio (SNR) than EPI data, as they are acquired in shorter acquisition windows, and at lower bandwidths. The shorter data acquisition window also leads to fewer issues with phase wrapping. Thus, B_0_ maps theoretically should provide a more robust base from which a warp field can be calculated. In practice, however, several factors from the present study suggest that reverse-blip correction may be generally more effective for FMRI studies.

First, with the population in this study (toddlers), acquiring reverse-blipped EPI data was more reliable because it could be acquired more quickly. Reverse-blipped EPI data typically require a few (i.e. approximately 15) volumes of an acquisition whose prescription matches the main EPI data set, meaning that these data can be acquired in under 30 seconds. The B_0_ field map data requires more time to acquire (approximately two minutes for this study). With a highly motion-prone population like the one scanned in this study, it is less likely to be collected, and it also has a higher chance of being corrupted with motion artifacts to the point of unusability. The speed with which reverse blip EPI data can be acquired (and reacquired, if a subject moves), makes it more suitable for distortion correction when scanning highly motion-prone populations.

Second, the acoustic noise generated by the B_0_ mapping can also cause issues, again given the present population and the conditions under which the study was conducted (late night/early morning sessions - when subjects would typically be asleep). The noise generated by the B_0_ mapping sequence is noticeably different from that generated by the EPI sequence used for the acquisition of the task or resting state data. This, in conjunction with it being a separate acquisition, makes it more likely to be disruptive and to awaken a sleeping subject. For the reversed-blip EPI correction, the difference in the noise characteristics is much less, and in most cases, indiscernible from the main acquisition. It may also be possible to integrate the reverse blip EPI scan as part of the main acquisition or its prescan portion, so collecting these data are better integrated into a scanning work-flow.

Third, the reversed-blip EPI data collection has advantages in its relative ease of acquisition. Because parameters are almost identical between the reversed-blip data set and the EPI data they are used to correct, there are fewer independent parameters that need to be optimized or recorded for these data to be used. For B_0_ field map data, echo spacings (both for the B_0_ field map and the EPI data) must be recorded and utilized in the various correction streams. The phase-encode direction of the EPI data must also be noted. Many of these parameters are not stored or recorded by scanner manufacturers in standardized ways, which can make implementing standardized correction pipelines more difficult, especially for cross-site and cross-scanner studies.

Fourth, the reverse-blipped EPI has greater similarity of the sequences used to generate the respective types of warp fields than B_0_ does. For the latter, commonly 2D gradient-echo based pulse sequences are used, and such pulse sequences have different characteristics and behavior than the EPI pulse sequences typically used for task and resting FMRI experiments. Most fundamental to these differences are the gradient waveforms used for each kind of acquisition; the eddy eddy currents generated by each kind of waveform; and the differing effects those eddy currents have on image distortion. Thus, even though the B_0_ is theoretically a direct measure of inhomogeneity, the waveform difference means that creating a warp field from information generated by the same sequence that will be used for imaging should be more representative, and give a warp field that is more representative of all of the effects causing image distortions. Besides favoring a warp field computed from reversed-blip data, this issue could be also be addressed by using EPI-based B_0_ field maps [11, 12].

Finally, while comparisons of cost functional results between B_0_ and reversed-blip techniques did not show statistical differences, direct structural comparisons showed that the latter performed at least as well as the former. Indeed, in several cases the outer gray matter boundaries appeared to be more aligned with structure using reversed-blip corrections (see subjects B, C and F in Fig. 2); the same was true for internal structures in many cases (e.g. subjects A, C, D and E in Fig. 2). These differences may be due to relative sequence characteristics, as noted above, as well as to the need for regularization, when using B_0_ data for correction.

Given all of these factors, acquisition and use of reverse-blipped EPI data seems to be a more robust and flexible option, given the relative speed with which it can be acquired, and its relative ease of use, compared to acquiring and using B_0_ data. Using B_0_-based techniques may be more appropriate in experiments where SNR is more constrained.

Another group [17] recently performed a similar comparison of multiple EPI distortion correction algorithms in high field (7T) animal data. That study used a smaller sample size (8 mice), but produced a more uniform set of data from all animals. From the multiple metrics computed and compared by the authors (Jacquard index, Cross Correlation, and Mutual Information), the authors determined that their implementation of the reversed blip correction also gave better results when compared with using a B_0_-field map based correction.

EPI sequences are commonly utilized for other types of MRI data acquisition, with diffusion weighted imaging (DWI) being a common application. In this context as well, the major methods for dealing with EPI distortion are blip-reversed correction [3, 13] and B_0_ field map acquisition [11, 14]. Further complications with DWI, however, are the presence of additional distortions arising from the eddy currents induced by the diffusion-weighting gradients, and must be corrected in conjuction with the static B _0_ inhomogeneities. Similar to the FMRI case of EPI acquisition, studies beyond this one have shown the benefits, if not necessity, of performing some kind of distortion correction [15, 16, 17].

One key disadvantage of both corrective methods used here is the problem of “pile-up” or aggregation of image energy, which leads to bright spots. Solutions cannot adequately determine the true undistorted values because of inherent information loss, and the unlimited number of possible solutions to the signal redistribution problem. It is likely that to deal with this loss of image information fully, one would have to process the original k-space acquisitions. In the end, it must be noted that while beneficial distortion correction can be performed in post-acquisition processing, it cannot remove all distortion effects from the EPI data. Therefore, researchers should still aim to minimize distortions in their acquisition protocols, in addition to implementing and performing the algorithmic corrections described here in their post-processing streams.

For this study, quantifying image alignment improvement between the anatomical data and the distortion-corrected EPI data, versus uncorrected EPI data, proved particularly challenging. The previously mentioned study [17] used Jaccard indices, intensity-based Cross Correlation, and histogram-based Mutual Information metrics, all showing similar performance for all tested correction schemes in that study (except for non-linear registration - in which only Mutual Information differed). The metric eventually used here (Pearson Correlation), and some tested previously (Mutual Information [15,17]), were also observed to perform similarly. What was critical to the results presented here was the use of T_1_ data that had been skull-stripped. Reducing signal sources from the skull (by using skull-stripped anatomical data) improved correspondence between what was being computed statistically, and what was being observed visually. Regardless of the metric chosen for evaluation, the authors found it critical to thoroughly survey all data visually, and the results of processing these data, to ensure that reported results are indeed consistent with the characteristics of the data. In the single case in this study, where the Z-PC score decreased after distortion correction, visual inspection still shows better geometric matching of the corrected EPI compared to the uncorrected EPI, with the T_1_ data.

In conclusion, results presented here clearly indicate that data to perform geometric distortion correction should be collected and utilized for all studies utilizing the EPI family of pulse sequences. Geometrically correcting task or resting-state FMRI data, or DTI data, will improve the alignment of these data to anatomical data, and will improve localization of function and pathologies to appropriate areas or regions in the brain, which should benefit both individual and group studies. The additional data required for either technique tested here can be acquired efficiently, particularly the blip-reversed correction which requires less than 1 minute of time to be added to a scanning session. The correction techniques are widely available and easily implementable in common software packages (as shown in the publicly available code repository referred to in this work), and should be applied as part of any EPI data analysis.

## Acknowledgments

This study utilized the high-performance computational capabilities of the Biowulf Linux cluster at the National Institutes of Health (NIH), Bethesda, MD, USA (http://biowulf.nih.gov), and was supported by the NIMH and NINDS Intramural Research Programs (IRP) at the NIH.

## References

1. Matthews, Hampshire. Clinical Concepts Emerging from fMRI Functional Connectomics. Neuron, Vol 91 (3), 511 – 528 (2016).

2. Jezzard, Balaban. Correction for geometric distortion in echo planar images from B0 field variations. Magn Reson Med, Vol 34(1), 65 – 73 (1995).

3. Andersson, Skare, Ashburner. How to correct susceptibility distortions in spin-echo echo-planar images: application to diffusion tensor imaging. NeuroImage, Vol 20, 870 – 888 (2003).

4. Holland, Kuperman, Dale. Efficient correction of inhomogeneous static magnetic field-induced distortion in Echo Planar Imaging. NeuroImage, Vol 50, 175 – 183 (2010).

5. Cox. AFNI - Software for Analysis and Visualization of Functional Magnetic Resonance Neuroimages. Computers and Biomedical Research, Vol 29, 162 – 173 (1996).

6. Smith, Jenkinson, Woolrich, Beckmann, Behrens, Johansen-Berg, Bannister, De Luca, Drobnjak, Flitney, Niazy, Saunders, Vickers, Zhang, De Stefano, Brady, Matthews. Advances in functional and structural MR image analysis and implementation as FSL. NeuroImage, Vol 23(S1), 208 – 219 (2004).

7. Avants, Tustison, Stauffer, Song, Wu, Gee. The Insight ToolKit image registration framework. Frontier in Neuroinformatics, Vol 8:44, 2014.

8. Fischl. FreeSurfer. NeuroImage, Vol 62, 774 – 781 (2012). DOI: 10.1016/j.neuroimage.2012.01.021

9. Cox, Glen. Nonlinear Warping in AFNI. Poster presented at the 19th Annual Meeting of the Organization for Human Brain Mapping; Seattle, WA, USA (2013).

10. Gorgolewski et al. The brain imaging data structure, a format for organizing and describing outputs of neuroimaging experiments. Scientific Data, Vol 3, 160044, 2016. DOI: 10.1038/sdata.2016.44

11. Reber, Wong, Buxton, Frank. Correction of Off Resonance-Related Distortion in Echo-Planar Imaging Using EPI-Based Field Maps. Magn Reson Med, Vol 39, 328 – 330 (1998).

12. Dymerska, Poser, Bogner, Visser, Eckstein, Cardoso, Barth, Tratting, Robinson. Correcting Dynamic Distortions in 7T Echo Planar Imaging using a Jittered Echo Time Sequence. Magn Reson Med, Vol 76, 1388 – 1399 (2016) DOI: 10.1002/mrm.26018

13. Irfanoglu, Modi, Nayak, Hutchinson, Sarlls, Pierpaoli. DR-BUDDI (Diffeomorphic Registration for Blip-Up blip-Down Diffusion Imaging) method for correcting echo planar imaging distortions. NeuroImage, Vol 106, 284 – 299 (2015).

14. Lee, Lazar, Lee, Holden, Terasawa-Grilley, Alexander. Correction of B0 EPI Distortions in Diffusion Tensor Imaging and White Matter Tractography. Proceedings of the ISMRM 12th Annual Meeting, Kyoto, Japan, p 2172 (2004).

15. Hutton, Bork, Joesphs, Deichmann, Ashburner, Turner. Image Distortion Correction in fMRI: A Quantitative Evaluation. NeuroImage, Vol 16, 217–240 (2002).

16. Irfanoglu, Sarlls, Nayak, Pierpaoli. Evaluating corrections for Eddy-currents and other EPI distortions in diffusion MRI: methodology and a dataset for benchmarking. Magn Reson Med, Vol 81, 2774–2787 (2019). https://doi.org/10.1002/mrm.27577

17. Hong, To, Teh, Soh, Chuang. Evaluation of EPI distortion correction methods for quantitative MRI of the brain at high magnetic field. Magnetic Resonance Imaging, Vol 33, 1098–1105 2015. DOI: http://dx.doi.org/10.1016/j.mri.2015.06.010

